# Dissecting planar and vertical organiser signals in early chick neural development

**DOI:** 10.64898/2026.03.20.713209

**Authors:** Alexandra S. Neaverson, Benjamin J. Steventon

## Abstract

Early neural development involves a combination of planar signals from the vertebrate organiser and vertical signals from its derived structures, the prechordal plate and notochord. However, the relative contribution of each structure to neural development is not clear. Here, we isolate anterior tissues from the primitive streak at successively later stages of development, to identify the extent of patterning that can occur prior to, during, and after the formation of the organiser and its later derivatives. Our results show that acquisition of neural identity occurs gradually and that exposure to planar signals from the developing node is necessary for neural plate specification. We also show that planar node-derived signals are required for AP patterning in isolated anterior tissues and give evidence that early neural tissue is of anterior character which subsequently becomes caudalised by signals (in part) from the developing node. However, anterior neural identity is lost without long-term contact with vertical signals from the axial mesendoderm. These results reveal a previously unappreciated level of autonomy in anterior neural development in the absence of node derived tissues.

**Summary statement:** Culture of isolated anterior tissues from the chick embryo reveal the roles of planar and vertical organiser signals for neural specification and anteroposterior patterning and maintenance.

## Introduction

Neural development and the organiser have long been considered intimately linked, since the landmark discovery that a graft of the amphibian Spemann-Mangold organiser can induce the formation of a secondary axis, containing a fully-patterning nervous system composed of host cells (Spemann, 1921; Spemann and Mangold, 1924). Equivalent structures were then identified in other organisms, including Hensen’s node in birds and mammals, located at the top of the primitive streak (Waddington, 1932, 1933, 1934, 1936, 1937; Waddington and Schmidt, 1933; Beddington, 1994; Kinder *et al*., 2001; Robb and Tam, 2004), with similar inductive abilities. This led to the concept of “neural induction” (NI), in which the neural plate develops in response to instructive signals from the node. As body axis development is accompanied by the self-differentiation of organiser derivatives such as the prechordal plate and notochord, an open question is the degree to which organiser signals operate via planar inductive interactions, or through vertical signals from organiser derivatives at later stages of development.

There are multiple distinct processes in early neural development for which planar vs. vertical signals may play distinct and instructive roles, beginning with the initial specification of the neural plate. Specification assays involving explants of the prospective neural plate develop into *SOX2*^+^/*SOX1*^+^ neural (lens-like) tissue in culture (Trevers *et al*., 2018), demonstrating the degree of autonomous neural development that the epiblast can achieve when isolated from its endogenous signalling environment. Indeed, vertical signals from the hypoblast play a role in ‘priming’ the epiblast for further neural development, with FGF signals inducing the expression of pre-neural genes such as *SOX3* and *ERNI* (Streit *et al*., 2000). However, early neural plate explants are sensitive to exogenous epidermal-promoting BMPs until HH4, after which point neural development is stable even in the presence of BMPs (Wilson *et al*., 2000; Wilson and Edlund, 2001). In the context of a node graft, organiser-derived BMP antagonists such as Chordin are considered required but not sufficient for the commitment of naïve tissue to neural fate (Streit *et al*., 1998), since additional organiser-derived signals are required – which are yet to be fully characterised (Lu *et al*., 2025). In the embryo, *SOX2* expression begins at HH4^+^, around the same time that neural plate specification is complete (Uwanogho *et al*., 1995; Rex *et al*., 1997; Linker and Stern, 2004; reviewed in Stern, 2005), and a node graft into the area opaca can induce the expression of *SOX2* (Streit and Stern, 1999; Streit *et al*., 2000; Linker and Stern, 2004; Trevers *et al*., 2023). It has not yet been tested whether the intact neural plate has a continued requirement for organiser-derived signals (including BMP antagonists) in order to acquire *SOX2* expression. Assessing whether planar organiser-derived signals are required for neural specification requires removing all sources of these signals in the embryo before node derivatives emerge, in addition to other primitive streak regions that are known to contribute to its regeneration (Waddington, 1932; Grabowski, 1956; Yuan, Darnell and Schoenwolf, 1995a, 1995b; Psychoyos and Stern, 1996; Joubin and Stern, 1999).

Following neural specification, the neural plate becomes subdivided into the prospective forebrain, midbrain, hindbrain and spinal cord territories (Storey, Crossley and Stern, 1992; Muhr *et al*., 1999; Wittler and Kessel, 2004; reviewed in Stern, 2001). A node graft can induce the entire spectrum of AP fates after grafting (Storey, Crossley and Stern, 1992; see also textbooks such as Gilbert and Barresi, 2017), but it is not clear the extent to which this depends on early planar signals or later vertical signalling. The derivatives of the organiser – the axial mesendoderm, made up of prechordal mesendoderm and the notochord – retain the ability to specify defined regions of neural tube along its AP axis (Izpisúa-Belmonte *et al*., 1993; Foley, Storey and Stern, 1997; Pera and Kessel, 1997; Rowan, Stern and Storey, 1999), suggesting the continued involvement of vertical signals acting from the mesendoderm to the overlying neuroepithelium. Indeed, studies in mice emphasise the importance of signals from the anterior axial mesendoderm for patterning and/or maintaining anterior neural tissues (Ang and Rossant, 1993; Camus *et al*., 2000; Martinez Barbera *et al*., 2000; Andersson *et al*., 2006; Fossat *et al*., 2015; Tam and Masamsetti, 2025). However, in the chick, removing the prechordal mesendoderm does not disrupt coarse AP patterning (Pera and Kessel, 1997), with the exception of disrupting the specification of hypothalamic tissue (reviewed in Manning and Placzek, 2024). Prechordal mesendoderm progenitors are located within a defined anterior compartment within the node (Kanno, Rothstein and Simoes-Costa, 2025); it is possible that much of their role in anterior patterning is complete before they begin their exit from the node at HH4^+^. In addition, signals from non-node sources, such as Wnt, FGF and retinoic acid, are thought to be involved in neural AP pattern establishment (Blumberg *et al*., 1997; Muhr, Jessell and Edlund, 1997; Gould, Itasaki and Krumlauf, 1998; Muhr *et al*., 1999; Wittler and Kessel, 2004; Nordström *et al*., 2006). It is not clear to what extent vertical signals from axial mesendoderm, planar signals from the organiser, and signals from non-axial sources all contribute to the formation and maintenance of the AP pattern.

During axial elongation, axial and paraxial tissues develop in close proximity, so it is possible that they depend on each other for proper morphogenesis, as they do in the tailbud (Xiong *et al*., 2020). In support of this, grafts of the prechordal plate have been shown to promote convergence of the overlying neuroepithelium (Yoshihi *et al*., 2022). In contrast, explanted pieces of the neural plate do not show signs of morphogenesis (Muhr, Jessell and Edlund, 1997; Muhr *et al*., 1999; Wilson *et al*., 2000, 2001; Nordström *et al*., 2006). Yet, *Xenopus* prospective neural plate explants can elongate when sandwiched together with an explant of the chordamesoderm or another neural explant, suggesting that basal contact with another tissue is required for neural plate morphogenesis. Therefore, a final open question is whether later stages of neural development can occur in the absence of underlying axial mesendoderm in the chick embryo.

Here we employ an anterior tissue isolation method to determine the role of planar signals from the organiser, compared to vertical signals from its derivatives, for neural specification and anterior-posterior patterning. We find that neural specification is a gradual process that requires planar signals from the anterior primitive streak, and that subsequent planar signals from the node act to quickly posteriorize this tissue. Despite anterior neural fates being established early, vertical signals from the underlying axial mesendoderm are important for maintaining forebrain identity. We also find a significant level of autonomy in morphogenesis; once established, the neural plate is able to form complex structures in the absence of other midline tissues.

## Results

### Planar signals from the developing organiser specify the neural plate

In normal development, *SOX2* is switched on in the early neural plate at HH4^+^ and is generally considered to be the earliest marker of the definitive neural plate (Supplementary Figure 4C; see also Uwanogho *et al*., 1995; Rex *et al*., 1997; Chapman *et al*., 2003; Stern, 2005; Sanchez-Arrones *et al*., 2012). Control of *SOX2* onset is a multi-step process that is not fully understood (Papanayotou *et al*., 2008). Since the organiser produces a wide range of secreted molecules in the period before *SOX2* expression is initiated, it is possible that this transition requires organiser-derived signals. However, it is also possible that earlier signalling activity from the developing primitive streak establishes commitment to later *SOX2* expression, or that organiser-derived signals are simply not required for this transition to occur.

Previous experiments (Gallera and Nicolet, 1969) and our own data (Supplementary Figure 1) indicate that the region of the embryo with organiser ability (i.e., the ability to induce neural gene expression in ectopic locations after grafting) is larger than the node itself. This is reflected in the expression of organiser-associated genes, some of which extend to roughly midway down the primitive streak along its AP axis (Supplementary Figure 2; see also Izpisúa-Belmonte *et al*., 1993; Chapman *et al*., 2002). Indeed, in previous node ablation experiments where the objective was to remove all notochord progenitors (which contribute to the node’s NI abilities (Kanno, Rothstein and Simoes-Costa, 2025)), a much larger area is excised (Psychoyos and Stern, 1996). Furthermore, excision of this region is followed by complete regeneration (Psychoyos and Stern, 1996; Joubin and Stern, 1999), so to study the timing of neural fate establishment in the absence of further node-derived signals, this regeneration must be prevented. We have used a simple method that we refer to as “anterior segment culture” (Figure 1A), in which the anterior part of the embryo (containing the prospective neural plate) is isolated from the primitive streak and posterior regions entirely. This is a variation on previous culture techniques (such as those used by Spratt Jr. and Haas, 1960; Lipton and Jacobson, 1974; Darnell, Stark and Schoenwolf, 1999, and Chapman *et al*., 2003), modified to ensure that all prospective neural plate regions were included. Confining the culture is necessary, to prevent the epibolic expansion of the extraembryonic tissue, using a soldering iron to denature the vitelline membrane as described by (Serrano Nájera *et al*., 2025). This permits the assessment of neural development in the absence of the node and primitive streak.

**Figure 1.**
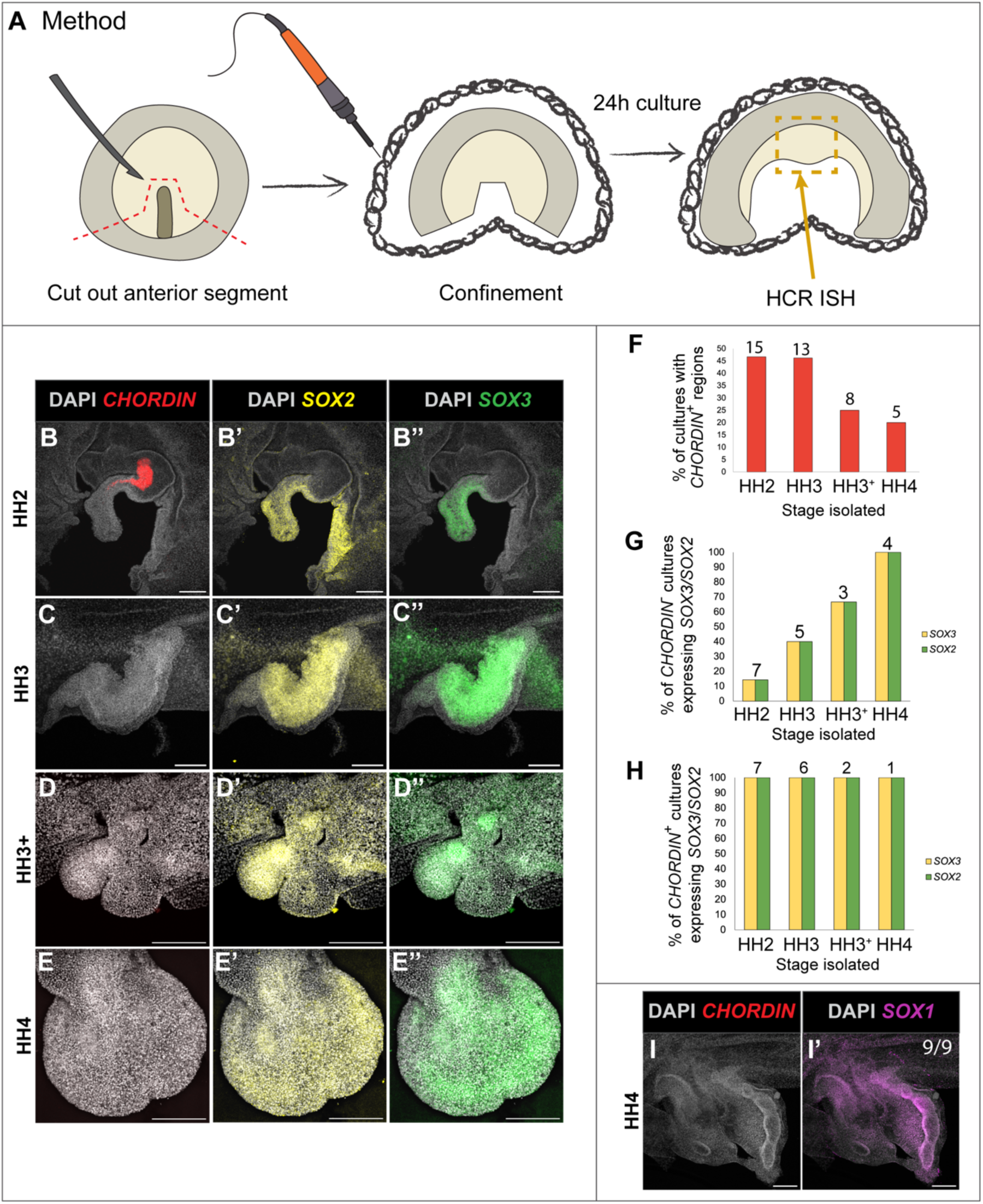
Neural specification occurs during gastrulation, as the node develops, and is reinforced by the node. (A) Anterior segments were created by cutting around the node and primitive streak at HH2, HH3, HH3^+^ or HH4. The segment was confined by burning a barrier in the vitelline membrane around the segment using a soldering iron. Segments were cultured for 24h before HCR FISH staining. (B-E’’) Expression of *CHORDIN*, *SOX2* and *SOX3* in the resulting neural tube-like protrusions from anterior segments after culturing for 24h. (F) Percentage of cultures with *CHORDIN*^+^ regions, indicating node regeneration. 7/15, 6/13, 2/8 and 1/5 cultures had *CHORDIN*^+^ regions at HH2, HH3, HH3^+^ and HH4 respectively. (G) Percentage of cultures that expressed *SOX3* and *SOX2* in the absence of *CHORDIN*. 1/7, 2/7, 2/3 and 4/4 cultures at HH2, HH3, HH3^+^ and HH4 respectively. (H) Percentage of cultures that expressed *SOX3* and *SOX2* that also had *CHORDIN*^+^ regions, showing that the presence of a *CHORDIN*^+^ region guarantees the expression of *SOX3* and *SOX2*. 7/7, 6/6, 2/2 and 1/1 cultures at HH2, HH3, HH3^+^ and HH4 respectively. (I-I’) Example of a HH4 anterior segment 24h after isolation, stained for *CHORDIN* and *SOX1*. (B-C’’) and (I-I’) are 5X maximum intensity projections, scale bars 200µm. (D-E’’) are 10X maximum intensity projections, scale bars 200µm. Numbers above bars on (F), (G) and (H) indicate sample size.

Anterior segments were isolated from embryos at HH2, HH3, HH3^+^, HH4, HH5 and HH6, and cultured for 24h. During this time, the early neural plate will sometimes autonomously develop backward-facing protrusions. The frequency of protrusion formation after isolation at each stage was quantified; isolation after HH4 was associated with an increased ability to form neural tube-like protrusions (Supplementary Table 1). HCR FISH was then used to visualise the expression of *CHORDIN*, *SOX2* and the pre-neural marker *SOX3* (Trevers *et al*., 2018, 2023) (Figure 1B-E’’). An assessment of *CHORDIN* expression in control embryos (Supplementary Figure 3) was used to inform the cutting position to minimise the likelihood of leaving behind *CHORDIN*^+^ primitive streak tissue, which may have NI (neural induction) ability.

In this assay, the indicator of neural specification is the ability to express *SOX2* in the absence of *CHORDIN*. Both HH2 and HH3 anterior segments frequently developed *CHORDIN*^+^ regions (Figure 1B, F) meaning that at these stages, it is difficult to dissociate neural specification from the potential inductive signals provided by the *CHORDIN*-expressing cells. The appearance of *CHORDIN*^+^ regions indicates the regeneration of node tissue along the cut edge of the culture (similar to observations by Caldarelli *et al*., 2024). Node regeneration therefore occurs more frequently in anterior segments from younger embryos.

The incidence of *SOX2/SOX3* expression without associated *CHORDIN*^+^ tissue increased with isolation at later stages, and we observe a stepwise increase in the frequency of *SOX2/SOX3* expression between HH2 and HH4 (Figure 1G). A small number of embryos formed neural tube-like structures and expressed *SOX2/SOX3* after isolation at HH2-3 (Figure 1C-C’’). This suggests that neural specification is a gradual process, rather than a switch occurring at a defined stage, and that the acquisition of neural plate identity is initiated prior node formation and complete by HH4 (100% at HH4, Figure 1G). Neural specification cues are therefore initially provided by planar signals from the PS, which contains precursor cells that will later form the node (Izpisúa-Belmonte *et al*., 1993; Foley, Skromne and Stern, 2000; Streit *et al*., 2000; reviewed in (Stern, 2005)). These are then superseded by signals from the node proper as it develops, since cultures that were able to regenerate *CHORDIN* tissue (or had *CHORDIN*^+^ left behind) expressed *SOX2* and *SOX3* in 100% of cases (Figure 1H), suggesting that although the process is initiated by node precursors, the node proper provides robustness to the neural specification process.

In later development, *SOX2* is expressed other cell types, such as placodes and neural crest cells, which include cells that do not contribute to the nervous system (Streit, 2002; Roellig *et al*., 2017). On the other hand, *SOX1* expression begins later in development (HH8, Supplementary Figure 4C) and is exclusively expressed in the neural tube (Uchikawa *et al*., 2011). HH4 anterior segments cultured for 24h were HCR FISH stained for *SOX1* expression (Figure 1I-I’); 9/9 cultures expressed *SOX1* in the absence of *CHORDIN*, demonstrating the progression of neural development in the absence of the node and its derived tissues. This suggests that once neural identity is specified, development can continue without additional planar NI signals.

### Stepwise posteriorisation of the neural plate requires node-derived signals

We next sought to determine whether vertical signals (i.e., those derived from node-derived axial mesendoderm tissues) are required for AP patterning, or whether node-derived, planar signals are sufficient (Figure 2). The expression of the marker genes used to identify AP patterning is shown in Figure 2C. *OTX2* is expressed in the forebrain and midbrain, while *KROX-20* is expressed as two bands in rhombomere 3 and 5 of the hindbrain. The anterior segment culture method was used (Figure 2A), separating the early neural plate from the primitive streak and posterior body at HH3^+^, HH4, HH5 or HH6 – using HH3^+^ as the youngest stage. Following a 24h culture period, anterior segment cultures were HCR FISH stained for *OTX2*, *KROX-20* and *CHORDIN*. At HH3^+^, 40% of cultures expressed *OTX2*, but none expressed *KROX-20* (Figure 2D-F, O-P). This contrasts to cultures isolated at HH4 and HH5, of which 62/67% expressed *KROX-20*, and 100% expressed *OTX2* (Figure 2G-L, O-P). After isolation at HH6, 100% of cultures expressed *KROX-20* in a pattern resembling the rhombomeres of the hindbrain (Figure 2M-N, P). These results show that AP pattern acquisition primarily occurs through cues from planar, node-derived signals between HH3^+^ and HH4, and support the idea that the first neural tissue to develop is of anterior character (“Activation” – as shown by the *OTX2*^+^ cultures isolated at HH3^+^) of which some parts are then transformed into more posterior regions (“Transformation”) (Nieuwkoop, 1952).

**Figure 2.**
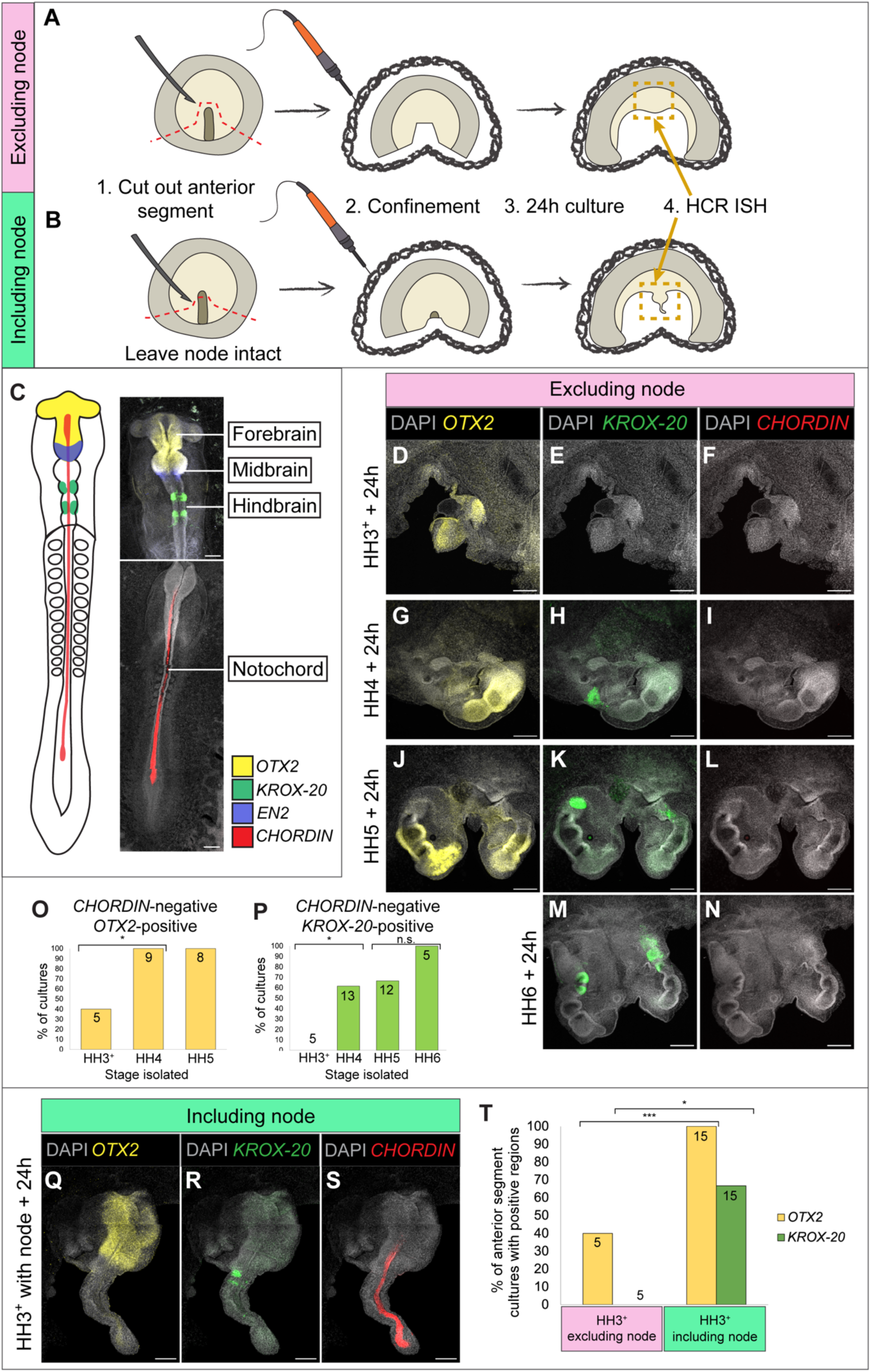
Anteroposterior patterning of the neural plate through planar signals from the node. (A) Diagram to show the method of creating and confining the anterior segment cultures. The primitive streak and surrounding area were cut away and removed, leaving the neural plate and lateral edges of the embryo with extraembryonic tissue attached, then confined as described in Figure 4. Anterior segments were cultured for 24h before HCR FISH staining. (B) The same process as in (A) was repeated, but the node was left intact. (C) Diagram to illustrate the normal expression pattern for AP patterning genes in the neural tube and *CHORDIN* expression in the notochord, alongside HCR FISH images. (D-N) DAPI staining and HCR FISH of anterior segments after 24h culture showing the expression of forebrain/midbrain (*OTX2*) and hindbrain (*KROX-20*) marker genes in the absence of node regeneration (*CHORDIN*). (O) Percentage of anterior segment cultures isolated from HH3^+^-HH5 embryos and cultured for 24h that expressed *OTX2*. Incidence of *OTX2* expression increases between HH3^+^ and HH4 (p = 0.027). (P) Percentage of anterior segment cultures isolated from HH3^+^-HH6 embryos that expressed *KROX- 20*. Incidence of *KROX-20* expression increases between HH3^+^ and HH4 (p=0.036). (Q-S) DAPI staining and HCR FISH of anterior segments isolated at HH3^+^ including the node, after 24h culture, showing the expression of *OTX2*, *KROX-20*, and *CHORDIN*. (T) Percentage of anterior segments isolated at HH3^+^ with or without including the node that expressed *OTX2* (yellow) and *KROX-20* (green). 2/5 expressed *OTX2* and 0/5 expressed *KROX-20*. With the node included, 15/15 expressed *OTX2* (p = 0.009) and 10/15 expressed *KROX-20* (p = 0.033). All statistical analysis was carried out using Fishers Exact test. Numbers overlaying bars indicate sample size. All images are 5X maximum intensity projections, scale bars 200µm.

The reduced expression of *KROX-20* relative to *OTX2* after isolation at HH4-5 could indicate that posteriorisation of the neural plate requires patterning signals for a longer duration than more anterior regions. These cues may be planar signals from the node, vertical signals from the later-forming axial mesendoderm, or planar signals from other tissues with known patterning properties – like the posterior streak/epiblast and paraxial mesoderm (reviewed in Wilson and Houart, 2004) that are absent from the culture. To separate these possibilities, the anterior segment cultures were repeated at HH3^+^ with the node left intact (Figure 2B). 100% of these cultures expressed *OTX2*, while 60% expressed *KROX-20* (Figure 2Q-T). However, 100% of cultures isolated at HH6 without the node/axial tissues expressed *KROX-20* (Figure 2P). This suggests that signals from axial tissues are sufficient and required for the stabilisation of *OTX2* expression and do contribute to the stabilisation of hindbrain fate, but non-axial tissues providing caudalising signals likely act in conjunction with the node between HH4-6 to stabilise *KROX-20* pattern. Overall, this indicates that AP patterning occurs in a stepwise fashion and is predominantly orchestrated by planar node-derived signals, but posterior fates take longer to establish and are supported both by vertical signals from axial tissues and caudalising signals from non-axial tissues.

### Neural morphogenesis can occur without node-derived axial mesendoderm

Previous studies indicate the importance of interaction between the axial tissues – the mesendodermal head process and prechordal plate, with the overlying neural plate – during axial extension and head formation (Xiong *et al*., 2020; Yoshihi *et al*., 2022). Yet, the anterior segment cultures produce extensive neural tube-like structures, especially when isolated at HH4 or later (Figure 1-2), showcasing the intrinsic ability of the neural plate to fold up and undergo morphogenesis in the absence of the mesendodermal and paraxial tissues (Moury and Schoenwolf, 1995; Schoenwolf and Yuan, 1995; Smith and Schoenwolf, 1997; Lawson, Anderson and Schoenwolf, 2001).

However, the orientation of marker gene expression in the neural tube-like structures is often inverted relative to normal; for example, in Figure 2G-I and J-L, the expression of the *OTX2* appears in a more posterior position than *KROX-20*. To better understand this apparent axis inversion, a timelapse recording of an anterior segment culture from a HH4 embryos was obtained; stills are shown in Figure 3C-C’’ (Movie S2). Masks over the forebrain, midbrain and hindbrain regions were manually added to demonstrate the movement of the tissues over time, compared to an unoperated control embryo (Figure 3B-B’’, Movie S1). In the absence of the midline, the early neural plate is split instead of developing as one coherent structure. The two halves develop and form complex structures independently (as described by Darnell, Schoenwolf and Ordahl, (1992), including inflated brain vesicles with then push towards anteriorly as they grow and collide with the opposite side. As they continue to grow, the two sides push against each other and end up protruding backwards into the gap (Figure 3C’’), resulting in an inverted, inside-out brain-like structure developing.

**Figure 3.**
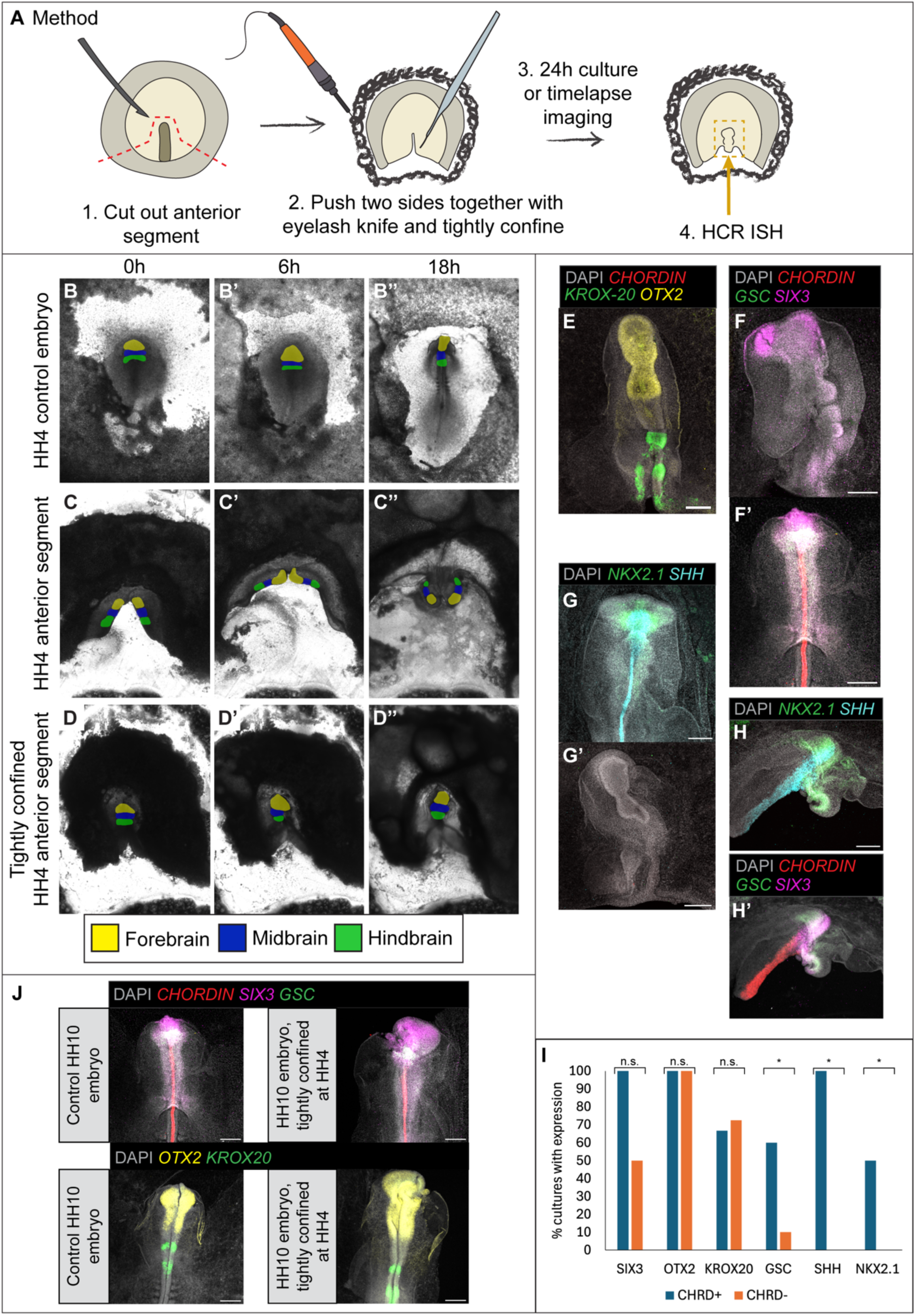
Anterior segment cultures become inverted without midline tissue, but can be rescued by tight confinement. (A) Diagram to show the method of tightly confining anterior segments. Method is as previously described in Figure 4, but additionally step 2 involves pushing the two lateral pieces of tissue together with an eyelash knife, and burning the barrier in the vitelline membrane more closely to the tissue. (B-D’’) Stills from timelapse movies showing the development of (B) a control embryo, (C) a HH4 anterior segment, and (D) a tightly confined anterior segment, during 18h culture. Masks were manually added to show the approximate position of the forebrain (yellow), midbrain (blue) and (hindbrain). (E, F) Tightly confined anterior segments after HCR FISH to demonstrate the correct relative positions of (E) *OTX2* and *KROX-20,* without *CHORDIN* present, and (F) *SIX3*, without *GSC* and *CHORDIN* present. (F’) Control HH10 embryo with HCR FISH staining for *CHORDIN*, *GSC,* and *SIX3*. (G) Control HH10 embryo with staining for *NKX2.1* and *SHH*. (G’) Tightly confined anterior segment with staining for *NKX2.1* and *SHH*. (H-H’) Tightly confined anterior segment demonstrating that *CHORDIN* expression is associated with the co-expression of *NKX2.1*, *SHH*, *GSC* and *SIX3*. (I) Quantification of the percentage of anterior segments (including both tightly and loosely confined samples) expressing markers of neural AP pattern (*SIX3*, *OTX2*, *KROX-20*), prechordal plate (*GSC*), floorplate/notochord (*SHH*), and ventral forebrain (*NKX2.1*), in the presence of absence of *CHORDIN*^+^ tissue. The presence of *GSC* (p = 0.042), *SHH* (p = 0.012) and *NKX2.1* (p = 0.039) are associated with the co-expression of *CHORDIN*. Total number of samples for each gene: For *CHRD*^-^samples, *SIX3* n = 20, *OTX2* n = 25, *KROX-20* n = 29, *GSC* = 20, *SHH* = 6, *NKX2.1* = 14. For *CHRD*^+^ samples, *SIX3* n = 5, *OTX2* n = 4, *KROX-20* n = 6, *GSC* = 5, *SHH* = 3, *NKX2.1* = 4. (J) HH10 control embryo (left) compared to a HH10 embryo tightly confined from HH4 (right) to demonstrate that tight confinement does not modify the overall localisation of expression for any of the genes tested. All statistical analysis was carried out using Fishers Exact test. All images are 5X maximum intensity projections, scale bars 200µm.

To understand whether this results from the altered geometry after removing the midline tissue, or whether the axial mesendoderm tissues are specifically required for orientation of the brain, the anterior segment culture method was modified slightly (Figure 3A). The two sides of the culture were forced into contact at the midline, then the culture was tightly confined to prevent expansion of the tissue and promote healing across the midline. Healing occurred and the neural plate was able to develop in this position rather than becoming inverted (Figure 3D-D’’, Movie S3). AP markers of the brain – *OTX2*, *KROX-20* and the anterior forebrain marker *SIX3* – were expressed in the correct orientation and position (Figure 3E, F; for *SIX3*, compare to normal expression in Figure 3F’), and the tissues developed into something that more closely resembles the normal brain. Tight boundaries for *OTX2* and *KROX-20* expression are observed (Figure 3E), reflecting the normal expression pattern (Figure 2C). Tight confinement itself was not found to impact the overall expression pattern for AP and axial mesendoderm markers compared to controls, when tested on whole embryos (Figure 3J). The frequency of *SIX3*, *OTX2* and *KROX-20* expression with and without *CHORDIN* present were compared, and no significant difference was found (Figure 3I). Overall, this shows that the AP orientation of the neural tube is stable once patterning has occurred (at HH4) so further signals from the axial mesendoderm do not substantially impact this AP pattern.

The remarkable AP patterning displayed in these cultures prompted us to ask whether they also showed signs of DV patterning. The axial tissues, including the prechordal plate, have a well-known role in specifying the DV axis of the brain by producing a graded SHH signal (Pera and Kessel, 1997; Briscoe and Ericson, 2001). However, SHH is produced by the node prior to the formation of the prechordal plate and notochord (Patten *et al*., 2003; Levine and Brivanlou, 2007). Furthermore, some evidence suggests that in the chick, specification of the floorplate – the ventral-most part of the neural tube – precedes the expression of SHH in the notochord (Kremnyov *et al*., 2018). Therefore, the first steps of DV patterning may begin prior to axial mesendoderm formation, through planar interactions. On that note, we looked for the expression of the floorplate/notochord marker *SHH*, prechordal plate marker *GSC*, and ventral forebrain/hypothalamic marker *NKX2.1* with HCR FISH in anterior segments isolated at HH4. *SHH* and *NKX2.1* expression were rarely observed (Figure 3G-G’); the few cultures expressing *NKX2.1* and *SHH* were also found to co-express *CHORDIN* (Figure 3H-H’), indicating that no ventral brain specification was observed in the absence of notochord tissue. In addition, the expression of *GSC* was also associated with the presence of *CHORDIN*, suggesting that prechordal plate tissue tends to be present in cultures with node regeneration or leftover node tissue. Signs of DV patterning only appear when *CHORDIN* is present, indicating that vertical signals from the axial mesendoderm are required for DV patterning, but not for AP patterning (Figure 3I).

### Long-term maintenance of forebrain identity requires node-derived axial mesendoderm

Anterior identity of neural tissues is established at or prior to HH3^+^, but studies in other vertebrate embryos, in particular the mouse, indicate that vertical signals from node-derived axial mesendoderm (e.g., the prechordal plate) are important for maintaining forebrain identity (Martinez Barbera *et al*., 2000; Andersson *et al*., 2006). If this holds true for the chick embryo, a decline in forebrain marker expression might occur following prolonged culture without node-derived tissues. Since *OTX2* is expressed in both the forebrain and midbrain, we additionally stained the cultures for *SIX3*, a forebrain-specific marker that initiates its expression at HH4 (Supplementary Figure 4B).

Anterior segments isolated at HH4 were cultured for 48h, before HCR staining for *OTX2* or *SIX3*, and comparing against anterior segments cultured for 24h (Figure 4). As previously described, after 24h culture, HH4 anterior segments consistently express *OTX2* (Figure 4A). At 48h, 4/6 *CHORDIN*-negative cultures expressed *OTX2*, but the fluorescence intensity was markedly reduced (Figure 4E). Most *CHORDIN*-negative cultures did not express *GSC* at 24h (Figure 4D) and none expressed *GSC* at 48h (Figure 4H), indicating that the prechordal plate was absent. For *SIX3*, expression at 24h appeared less frequently than *OTX2* – 10/25 cultures lacking *CHORDIN* expressed *SIX3* (Figure 4C). At 48h, 0/8 *CHORDIN*-negative cultures expressed *SIX3* (Figure 4G). Forebrain identity is therefore sensitive to signals from the axial mesendoderm when cultured for a prolonged period.

**Figure 4.**
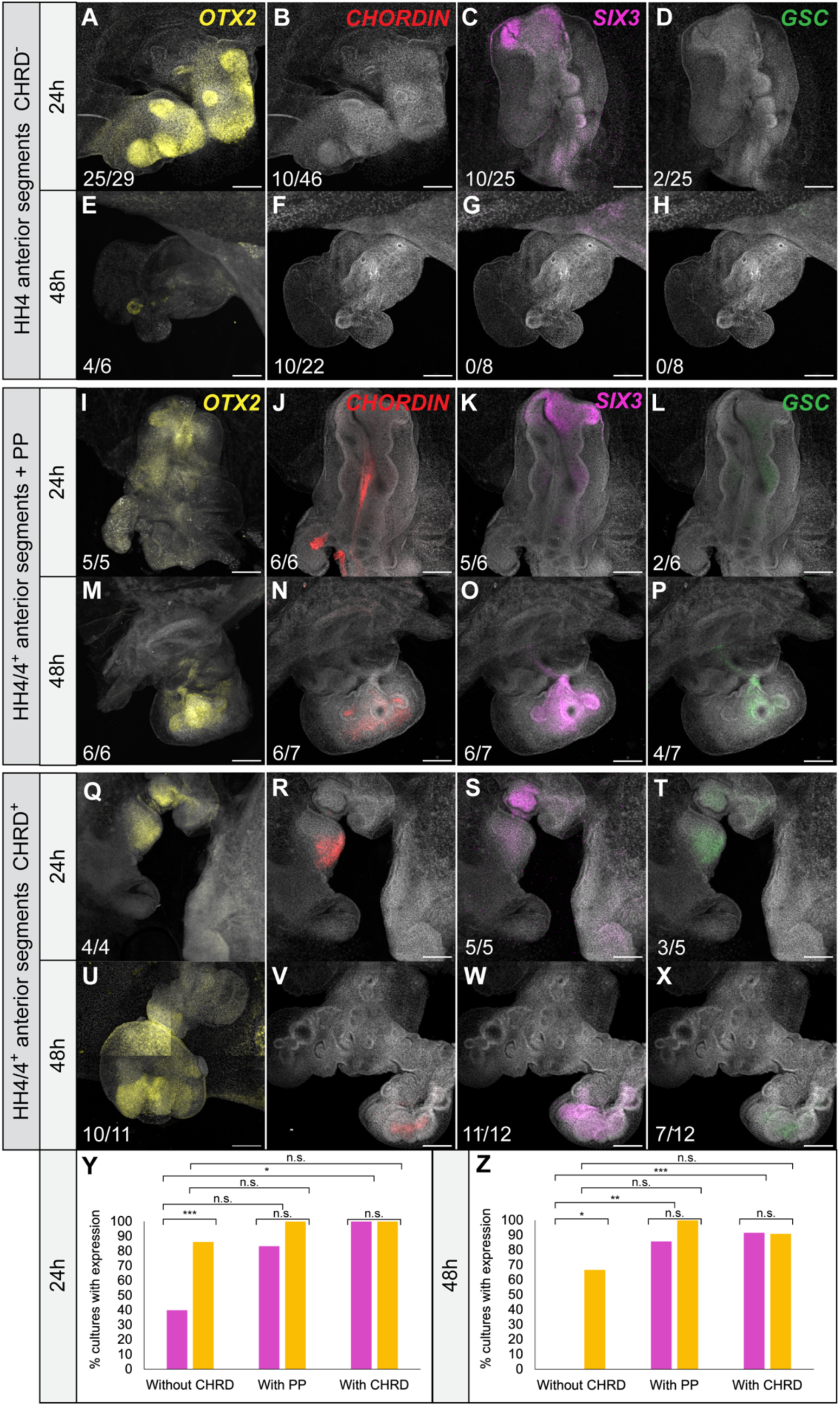
Loss of forebrain identity in the absence of the node and its derivatives. (A-H) Anterior segments were isolated from the primitive streak and posterior part of the embryo at HH4 as previously described, and cultured for 24h (A-D), or 48h (E-F) before HCR FISH staining for *OTX2*, *CHORDIN*, *SIX3* and *GSC*. Images and quantification are for samples that did not express *CHORDIN*. (I-P) Anterior segments were isolated from the primitive streak and posterior part of the embryo at HH4-4^+^, but with prechordal plate (PP) tissue left intact, then cultured for 24h (I-L) or 48h (M-P) before HCR FISH staining. (Q-X) Anterior segments were isolated from the primitive streak at HH4-4^+^ as previously described, and cultured for 24h (Q-T) or 48h (U-X) before HCR FISH staining. Images and quantification are for samples that did express *CHORDIN*. Scores indicate the number of samples positive for any given gene. All images are 5X maximum intensity projections, scale bars 200µm. (Y-Z) Quantification of the percentage of anterior segments at 24h (Y) or 48h (Z) with expression of *OTX2* (orange) and *SIX3* (magenta). At 24h (Y), without *CHORDIN* present*, OTX2* is expressed in significantly more samples than *SIX3* (p = 0.0005), and *CHORDIN* is associated with co-expression of *SIX3* (p = 0.042). At 48h (Z), *SIX3* is not expressed in any samples without *CHORDIN,* while *OTX2* is expressed in 67% of samples (p = 0.015). The presence of the prechordal plate (PP) (p = 0.0014) or *CHORDIN* expression (p = 0.00007) are associated with the expression of *SIX3* at 48h.

To test whether the reduced expression of *SIX3* could be rescued by the prechordal plate, the cultures were repeated at HH4-HH4^+^ with this tissue left intact (Figure 3I-P). At HH4^+^, the prechordal plate can be seen as the first cells begin to migrate anteriorly out of the node, while at HH4, these cells are still contained within the node in its anterior-most quadrant (Kanno, Rothstein and Simoes-Costa, 2025). We could visualise *GSC* expression in only 2/6 cultures at 24h, and 4/7 cultures at 48h likely due to a combination of weak expression, rare cell population and depth of the cells within the sample. However, the presence of axial mesendoderm was confirmed by the presence of *CHORDIN* (Figure 3J and N). Prechordal tissues did not significantly impact the expression of *SIX3* or *OTX2* at 24h, but boosted the expression of *SIX3* at 48h (Figure 4O, Y-Z).

Several cultures included *CHORDIN*^+^ tissue, either as a result of node regeneration or incomplete removal of node tissues. All cultures with visible *CHORDIN* (including those with the prechordal plate included) were grouped and compared to those without *CHORDIN* (Figure 4Q-X, Y-Z). At 24h, the presence of *CHORDIN* was associated with the presence of *SIX3*. By 48h, this association strengthens, as *SIX3* was not found in any cultures lacking *CHORDIN*. No significant association was found for *OTX2*.

Overall, this suggests that prolonged contact with signals from the underlying axial mesendoderm is important for long-term expression of forebrain-specific genes. This is particularly apparent for forebrain-specific *SIX3*, while *OTX2* – which could indicate forebrain or midbrain tissue – is retained but appears to be only weakly expressed. Since isolation of these anterior cultures was performed at HH4-4^+^ (when *SIX3* can be first detected in the anterior neural plate), from these findings we cannot rule out that some cultures may have been isolated before the onset of *SIX3* expression. These findings suggest a late role for the node, or a role for its early axial derivatives, in the initial induction and/or maintenance of *SIX3* expression.

## Discussion

In this work, we separate the involvement of planar and vertical signals during early chick development, providing a timeline for the involvement of the developing organiser, the node proper, the node-derived axial mesendoderm for the development and maintenance of neural tissues in the chick embryo. We propose a model for the involvement of these tissues in Figure 5.

**Figure 5.**
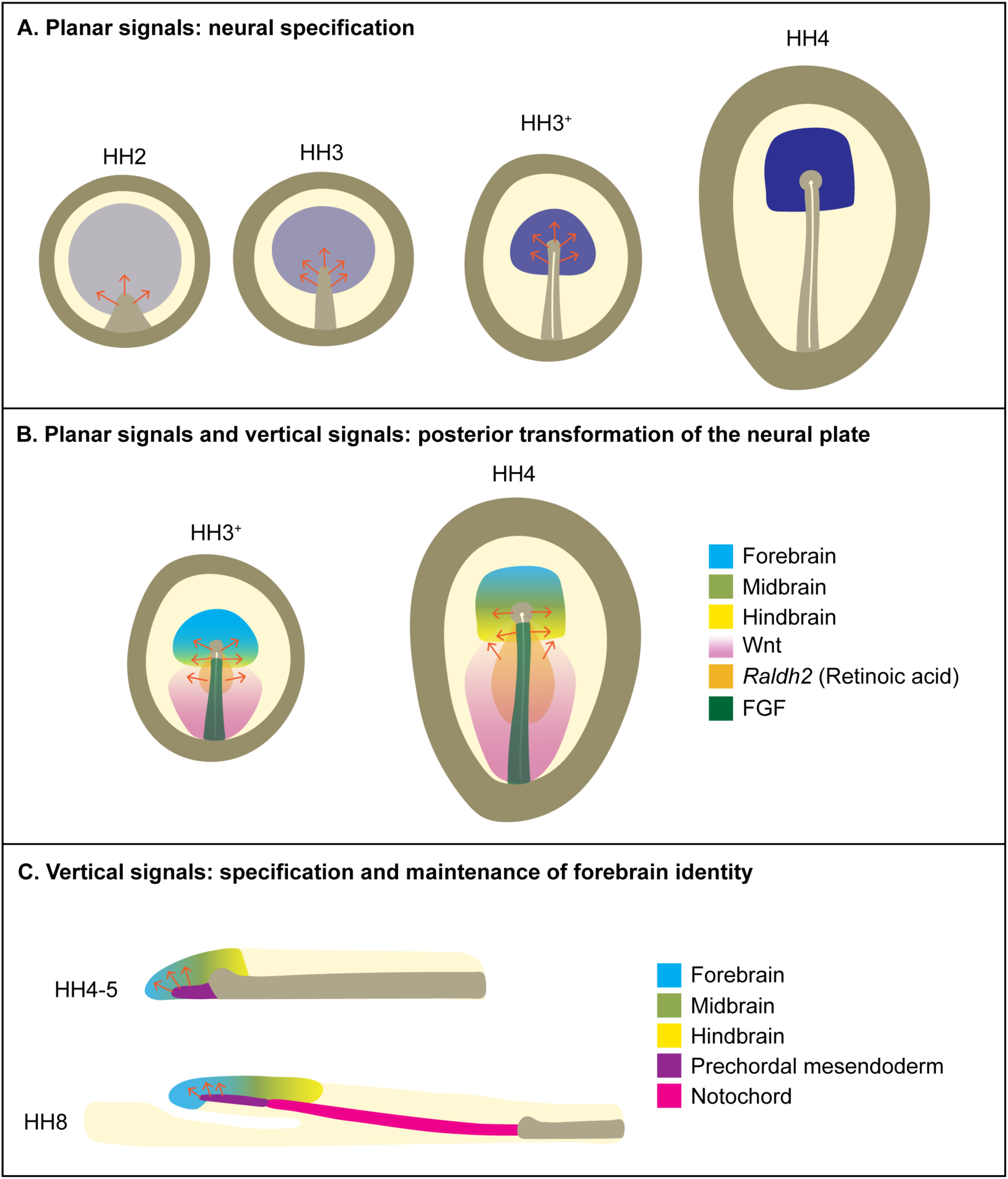
Summary of the contribution of planar and vertical signals to neural specification, AP regionalisation and forebrain patterning in early chick development. **(A) Planar signals act to specify the neural plate.** By HH2, the epiblast exists in a pre-neural, forebrain-like state, primed for neural specification. During HH2-3, the primitive streak develops, providing signals that pass along the plane of the epiblast to specify the neural plate. This is a gradual process that occurs concomitantly with primitive streak and node development. It continues through HH3^+^, when the neural specification signals become concentrated in the anterior half of the primitive streak including in the developing node. By HH4, the neural plate is now specified and *SOX2* expression begins. **(B) Planar and vertical signals contribute to the posterior transformation of the neural plate.** As the neural plate is becoming specified, it also becomes regionalised along its AP axis. The node provides planar signals that act to transform adjacent parts of the neural plate (previously of anterior character) into posterior neural tissue, specifying the midbrain and hindbrain by HH4. Planar posteriorising signals are also provided by the primitive streak (FGF) and the posterior epiblast (Wnt). At the same time, paraxial mesoderm progenitors ingress through the primitive streak and start to emerge beneath the epiblast, expressing *RALDH2* to provide retinoic acid signals that also contribute to the neural plate’s posterior transformation. **(C) Vertical signals contribute to the specification and maintenance of forebrain identity.** At HH4-5, prechordal mesendoderm (including prechordal plate) cells migrate anteriorly out of the node, providing signals that act vertically on the overlying neural tissue to specify the anterior/ventral forebrain. During the next 24h of development, the prechordal mesendoderm and notochord have become resolved and continue to provide vertical signals that set up the DV axis of the neural tube, and the prechordal mesendoderm continues to signal to maintain and refine neural identities in the anterior/ventral forebrain.

The anterior segment culture method permits the separation of the neural plate from sources of NI signals – allowing us to exclude the entire primitive streak – at defined stages. In doing this, we show that, rather than a simple switch, neural fate is gradually established through planar signals from the primitive streak during node development (Figure 1). 40% of anterior segments isolated at HH3 were able to achieve *SOX2* expression after 24h culture (Figure 1G). Small neural plate explants from HH3 embryos develop into *SOX2*^+^ neural tissue in culture, but upon exposure to BMP signals develop into epidermis (Muhr *et al*., 1999; Wilson and Edlund, 2001). Then, at HH4, a change occurs that allows neural plate explants to develop into neural tissue even in the presence of BMP. We show that our cultures, which can be thought of as whole neural plate explants, rely on organiser-derived signals between HH2 and HH4 in order to develop *SOX2*^+^ neural tissue. Crucially, our experiments demonstrate that the neural plate without an intact organiser will not initiate *SOX2* expression, and that *SOX3* expression – although initiated before primitive streak development (Rex *et al*., 1997) – drops unless neural specification has occurred, since *SOX3* is never observed in the absence of *SOX2*. This fits with the observations from organiser graft experiments showing that the initial ‘pre-neural’ state of the epiblast, induced by FGF, is unstable unless exposed to further signals from the organiser (Streit *et al*., 2000). Our results show that signals from the organiser (defined as the anterior half of the primitive streak at HH2-HH4) are required for *SOX2* expression and neural specification during normal development.

Anterior segments separated from organiser-derived signals at HH4 continue to develop and initiate expression of *SOX1* (Figure 1I-I’), a pan-neural gene that is not expressed until HH8 (Supplementary Figure 4D). This suggests that once the neural plate is specified, organiser-derived signals are no longer required, and neural development can progress autonomously. This may be the result of an auto/cross-regulatory loop involving the *SOXB1* genes (which include *SOX1*, *SOX2* and *SOX3*), since ectopic expression of *SOX2* has been shown to cause *SOX1* expression (Graham *et al*., 2003). Therefore, *SOX2* may directly initiate *SOX1* expression.

The anterior segment culture experiments are similar to the rostral blastoderm isolates performed by (Darnell, Stark and Schoenwolf, 1999) and (Chapman *et al*., 2003). They found that neural specification occurs by stage 3d (which corresponds to between HH3^+^ and HH4) which broadly fits with our timeline. A key distinction between our work and previous experiments is the use of multiplex HCR FISH to simultaneously stain for organiser genes and neural plate markers. This was important, since we frequently detected regions of *CHORDIN* expression indicating node regeneration. It allowed us to conclude that the presence of *SOX3/SOX2* in cultures isolated at early stages (e.g., HH2) is often correlated with the presence of a regenerated node, and to therefore exclude these samples from quantification. Furthermore, the anterior segments were grown in EC culture (as opposed to Spratt’s technique (Spratt, 1947)), which likely permits morphogenesis in a way that more closely resembles normal development, since the tissues remain in contact with the vitelline membrane. Our anterior segment cultures are somewhat similar to the classical ‘animal cap assays’ performed in *Xenopus*, which involve culturing the animal cap, previously exposed to early neuralising signals (Stern, 2005) in isolation or in the presence of other tissues or defined signalling molecules (Green, 1999). However, our cultures include extraembryonic tissue, a known source of anti-neural BMP signals (Streit *et al*., 1998). This allowed us to test the requirement of organiser-derived signals in a system that more closely resembles the normal signalling environment of the developing neural plate.

Previous research across vertebrate models broadly demonstrates that AP patterning can occur in the absence of vertical signalling from the underlying organiser-derived mesendoderm (Doniach, Phillips and Gerhart, 1992; Pera and Kessel, 1997; Schier *et al*., 1997; Strähle *et al*., 1997; Feldman *et al*., 1998; Fossat *et al*., 2015; Kakebeen *et al*., 2021; Manning and Placzek, 2024). However, prechordal plate removal experiments do not exclude the possibility that AP patterning signals are produced by these cells as they migrate out of the node. To exclude this possibility, experiments that completely prevent the formation of anterior axial mesendoderm are needed; this is achieved when anterior segment cultures are performed in embryos at HH4 or younger. The results of our anterior segment experiments stained for *OTX2* and *KROX-20* (Figure 2) are concordant with previous research: defined forebrain/midbrain and hindbrain tissue emerge within 24h, after isolation from axial mesendoderm before any vertical signalling could have occurred. These experiments additionally reveal differences in the timing of signal requirement for the anterior neural tissues compared to the more posterior neural tissues. We see that some anterior segments isolated at HH3^+^ can maintain the expression of *OTX2*, and this increases to 100% at HH4 (Figure 2O). Conversely, anterior segments isolated at HH3^+^ are not able to express *KROX-20*; this increases to 67% of cultures at HH4, then further increases to 100% of cultures at HH6 (Figure 2P). This suggests that genes expressed in the early embryo that later become restricted to the anterior – like *OTX2* – become committed before genes expressed in more posterior neural tissues, like *KROX-20*, giving evidence for Nieuwkoop’s Activation-Transformation model of neural development (Nieuwkoop, 1952). The increasing incidence of *KROX-20* expression up to HH6 suggests that additional signals and/or time are required to specify the hindbrain compared to the fore-/midbrain. The lack of sharp boundaries of *KROX-20* expression in culture isolated prior to HH6 could reflect insufficient contact with hindbrain-promoting Wnt/RA signals before isolation, or contributions from the underlying axial mesoderm to refining AP pattern (Rowan, Stern and Storey, 1999). Overall, our findings on AP patterning align with previous experiments using neural plate explants, showing the stepwise posteriorisation of the neural plate over time (Muhr, Jessell and Edlund, 1997; Muhr *et al*., 1999; Nordström, Jessell and Edlund, 2002; Nordström *et al*., 2006), and show that their findings extend to a whole-neural plate context.

In Figure 3, we demonstrate that early brain orientation and morphogenesis does not depend on the presence of underlying axial mesendoderm. The continued morphogenesis is consistent with experiments showing that neural tube elongation continues when notochord extension is stalled (Charrier *et al*., 1999) or when the notochord is absent in mouse mutants (Ang and Rossant, 1993). These results are similar to those of previous tissue isolation experiments (Smith and Schoenwolf, 1991; Moury and Schoenwolf, 1995; Schoenwolf and Yuan, 1995; Lawson, Anderson and Schoenwolf, 2001), further confirming that morphogenetic forces in anterior neural tissues are tissue-intrinsic.

Despite the strong resemblance to normal brain development, we show that anterior segments isolated at HH4 do not have any signs of patterning along the DV axis, as shown by the lack of *SHH* and *NKX2.1* expression (Figure 3G’). The only cultures expressing these genes were those that also expressed *CHORDIN*, indicating the presence of notochord tissue, which has a known role in DV patterning (Placzek *et al*., 1990; Yamada *et al*., 1991). A gradient of secreted SHH from the notochord and floorplate is the primary driver of DV patterning (Echelard *et al*., 1993; Roelink *et al*., 1994; Ericson *et al*., 1995, 1996). Although SHH is expressed in the node prior to the extension of the axial mesendoderm, our results suggest that the patterning of the neural tube’s DV axis does not occur until after these mesendodermal tissues come to underlie the developing neural tube. In contrast to AP patterning, which is well underway by HH4, DV patterning occurs later through vertical signalling.

Finally, extended culture of anterior segments revealed that genes associated with forebrain/midbrain identity are not maintained in the absence of axial mesendoderm (Figure 4). The fading expression of *OTX2* at 48h (Figure 4E) resembles effects seen when the early emerging prechordal mesendoderm is removed at HH5 (Pera and Kessel, 1997). The effect on *SIX3* was more pronounced, with fewer than half of the cultures expressing *SIX3* at 24h (Figure 4C). Since *SIX3* expression begins at HH4 (Supplementary Figure 4B), this can be interpreted in two ways: either its expression has already begun to decline at 24h, or the isolation was performed before the onset of expression, and *SIX3* was never switched on in a subset of the cultures. In either case, this points to an extended role for the node or its derivatives in promoting the expression of forebrain-specific genes. This likely includes a prolonged maintenance role for the axial mesendoderm, since *SIX3* is only maintained for 48h in cultures with *CHORDIN* or prechordal plate tissue present (Figure 4K, O, S, W). Similarly, *Gdf1*^-/-^ *Nodal*^+/-^ mouse embryos have impaired anterior axial mesendoderm formation, leading to a lack of *SIX3* expression and significant defects in the forebrain. These effects may stem from an excess of Wnt signalling, which is known to decrease *SIX3* expression (Lagutin *et al*., 2003). The node-derived prechordal mesendoderm likely provides signals that maintain forebrain identity, in part by providing Wnt antagonists (Foley, Storey and Stern, 1997; Rowan, Stern and Storey, 1999; Chapman *et al*., 2004; Fossat *et al*., 2015).

After the original organiser graft experiments by Spemann and Mangold (Spemann, 1921; Spemann and Mangold, 1924) a desire to find homologous structures, and the conflating of the results from graft experiments with normal development, has led to confusion surrounding the timing and mechanisms of neural induction (Martinez Arias and Steventon, 2018). Node graft assays have been useful for identifying the changes associated with neural induction and result in the development of a body axis (Dias and Schoenwolf, 1990; Storey *et al*., 1992; Trevers *et al*., 2018). However, they cannot necessarily be extrapolated to the node’s role in normal development. An emerging consensus view across vertebrates is that neural specification and regionalisation must occur earlier than morphological organiser development since ablation of the organiser (across vertebrate model organisms) does not significantly alter nervous system development and AP patterning (reviewed in Neaverson and Steventon, 2025). Using culture techniques with the neural tissue isolated from the organiser at progressively later stages, this allowed us to pinpoint the periods of signal requirement for different aspects of neural development. This allowed us to definitively show that signals from the node and its precursors are important for the onset of *SOX2* in the neural plate, and subsequent posteriorisation of the neural plate through planar signals. These are then supported by vertical signals from the emerging axial mesendoderm, which continue to provide local patterning signals that maintain AP pattern and induce DV pattern in the overlying neural tissue.

## Materials and Methods

### Ex ovo culture of chick embryos

Shaver brown chicken eggs were obtained from MedEggs (Henry Stewart & Company, Norfolk, England). Eggs were incubated at 37.5°C in a humidified incubator within 1-2 days of arrival. Incubation timings to achieve the desired stage were guided by Hamburger and Hamilton, (1992); for HH4, 18-22h of incubation was necessary. Embryos were cultured using the EC culture method (Chapman *et al*., 2001). Embryos in culture were imaged and staged using the Leica MZ10F stereoscope with brightfield illumination, typically at 2.5X unless otherwise specified. For transgenic GFP embryos or grafts, a separate image was obtained of the GFP fluorescence, which was then combined with the brightfield image using FIJI, to produce a single composite image.

### Node and streak grafting

Transgenic cytoplasmic GFP eggs from the Roslin Institute (University of Edinburgh, Easter Bush, Scotland) (McGrew *et al*., 2008) were incubated at 37.5°C for 15-17h to achieve HH3^+^ embryos to use as donors. Wild-type embryos incubated for the same duration were put into EC culture to be used as hosts. Donor GFP embryos were removed from the yolk and vitelline membrane and transferred into PBS. The region to be grafted was cut out using a microdissecting knife then aspirated into a 10µL pipette and transferred to the host embryo culture. Using an eyelash knife, a small ‘pocket’ or hole was created in the mesendoderm layer at the outermost edge of the area pellucida. The piece of tissue was then pushed into the hole with the mesendoderm side facing down. Excess saline was then removed using a pipette, before incubation at 37.5°C for 24h.

### Anterior segment cultures

Using HH2-HH6 embryos in EC culture, the primitive streak and posterior part of the embryo (including the associated area opaca tissue) were dissected out using a microdissecting knife. Cuts were made slightly anterior to the node, then parallel to the primitive streak on either side, stopping just beyond the region corresponding to the prospective spinal cord (guided by fate maps in Fernández-Garre *et al*., 2002). The segment was isolated by cutting diagonally across the embryo in a posterior-lateral direction, cutting through the area opaca, before removing the tissue with a pipette. Yolk and saline were removed from the embryo and the surrounding vitelline membrane. Then, a soldering iron set to 250°C was used to burn small regions of the vitelline membrane all around the embryo, creating a roughly circular barrier all around, to keep the anterior segment confined within the circle (Serrano Nájera *et al*., 2025).

### RNA Extraction

HH3+ embryos were dissected from eggs into cold PBS (Sigma-Aldrich). The primitive streak of each embryo was divided up into quarters, to give one node fragment (100% region), and one 75%, one 50% and one 25% region. 18 embryos were used in total, with corresponding streak sections pooled and transferred into 1mL Trizol (ThermoFisher). Tissues were homogenised with a handheld tissue homogeniser, then 0.2mL chloroform was added, before incubating at RT for 3 minutes. Samples were centrifuged for 15 minutes at 12,000 x g, 4°C. The upper aqueous phase was transferred to a new tube, then 0.5µL GlycoBlue and 0.5mL isopropanol was added, before incubating for 10 minutes at RT, then centrifuging for 10 minutes at 12,000 x g, 4°C. The supernatant was discarded and then pellet washed in 75% ethanol, then air dried at RT for 10 minutes. The pellet was then resuspended in nuclease-free water and incubated at 55°C for 15 minutes.

### RT-qPCR

Reverse transcription of RNA to cDNA was carried out using the Superscript III First-Strand Synthesis System (ThermoFisher) at 50°C for 45 minutes, before inactivating the reaction by heating to 70°C for 15 minutes. RNA was then removed using RNAse H digestion for 20 minutes at 37°C. qPCR was then performed using the QIAGEN Rotor-Gene PCR cycler and SYBR Green Mastermix, with reactions run in triplicate. Gene expression was quantified by standard curve analysis using a dilution series of cDNA from HH5 whole embryos. The Ct (threshold cycle) was calculated for each dilution in the series, using a threshold of 0.02, and used to create a log dilution-Ct graph for each gene. Using this, the relative concentration of each transcript in the unknown samples could be calculated. Dilutions were linearised and standardised relative to *ACTB*. Primer sequences can be found in Supplementary table 1.

### Embryo dissection, fixation and dehydration

Embryos were fixed by placing the embryo (from EC culture, attached to filter paper) in 4% PFA (paraformaldehyde, Sigma Aldrich, 158127) in PBS, overnight at 4°C. The following day, embryos were removed from the vitelline membrane using sharp forceps and transferred to PBST (PBS + 0.1% Tween). Embryos were then dehydrated using a series of 10 minute graded methanol/PBST washes, starting with 2 x PBST washes, then 25% methanol, 50%, 75%, then 100% methanol followed by storage at -20°C in 100% methanol.

### HCR RNA FISH

Gene expression was visualised using HCR RNA In Situ Hybridisation v3 (Choi *et al*., 2018). All buffers used for HCR were made in-house, according to the HCR™ RNA-FISH protocol (v3) for whole-mount chicken embryos. Embryos were first rehydrated using 10 minute graded methanol/PBST washes in reverse order. The PBST was removed and replaced with a 10 µg/mL solution of Proteinase K (Sigma, P4850) in PBST for 3 minutes at room temperature. This was replaced with 4% PFA for postfixing, for 20 minutes at room temperature, followed by 2 x washes with 5X SSCT (SSC buffer, Thermo Scientific, J60839.K2, + 0.1% Tween). Embryos were then incubated for 30 minutes in Hybridisation Buffer at 37°C. HCR probes (obtained from Molecular Instruments, information provided in Supplementary table 2, with the exception of the NKX2.1 probes, which were generously provided by Kavitha Chinnaiya, University of Sheffield) were diluted in Hybridisation Buffer to 4 pmol/mL and applied to the embryos for overnight incubation at 37°C. The next day, embryos were washed using 4 x 20 minute washes with Probe Wash Buffer at 37°C, then 2 x 5 minute washes with SSCT. Embryos were then incubated in Amplification Buffer for 5 minutes. Fluorescent HCR hairpins (Molecular Instruments) were snap-cooled by incubation at 95°C for 90 seconds before cooling to RT. The hairpins were diluted to 60 pmol/mL in Amplification Buffer. Pre-amplification solution was removed and replaced with hairpin solution, followed by overnight incubation at RT in the dark. Hairpin solution was removed, and embryos were washed 3 x 15 minutes with SSCT. DAPI staining was performed by incubating embryos in DAPI (1:1000 in SSCT) for 30 minutes at RT. DAPI solution was removed and embryos were washed for 3 x 10 minutes in SSCT.

### Confocal imaging

Fluorescently-stained embryos were mounted in VectaShield (Vector Laboratories, H-1000) between two glass coverslips (a smaller coverslip placed on top of a larger coverslip), separated by a layer of electrical tape. The edges of the small coverslip were sealed with nail polish. Confocal imaging was performed using the inverted Zeiss LSM700 microscope using either the 10X or 20X air objective.

### Image processing

Confocal images were processed in FIJI (Schindelin *et al*., 2012) to produce maximum intensity projections or summed slice projections. In some cases, individual images were stitched together using the ‘Pairwise Stitching’ function (Preibisch, Saalfeld and Tomancak, 2009). Images of embryos containing both a brightfield and a GFP channel were overlaid using the ‘Merge Channels’ function. The area of *SOX2* expression in Figure 1 was measured using the polygon tool and ‘Measure’ tool.

### Timelapse imaging

Live imaging of control embryos and anterior segments was performed using Nikon Eclipse Ti-E inverted widefield microscope with a heated chamber (37°C), using a 10X air objective. Images were obtained at 45 minute intervals. Movie files (.nd2) were processed in FIJI. For visualisation purposes, the forebrain, midbrain, and hindbrain regions were manually tracked from the brightfield movies and overlaid on the timelapse recordings.

### Wholemount imaging

Embryos were imaged using the Leica MZ10F stereoscope with brightfield illumination, typically at 2.5X unless otherwise specified. For embryos containing grafted regions, separate images were obtained of the GFP and brightfield channels, which were then combined in FIJI as previously described.

### Statistical analysis

To assess differences in gene expression between conditions, a Fisher’s Exact test was performed on 2×2 contingency tables comparing the number of embryos with positive and negative expression for each gene across conditions. Statistical analyses were carried out in Rstudio. A p-value threshold of 0.05 was used to determine statistical significance. Fisher’s Exact test was selected as it is appropriate for small sample sizes and computes exact rather than approximated p-values.

## Acknowledgements

We would like to thank the Steventon lab (especially Yuri Takahashi) for helpful comments and discussion surrounding the manuscript, Guillermo Serrano Nájera for assisting with creating the masks over the prospective brain regions in Movie S1-S3, Kavitha Chinnaiya and Marysia Placzek for the NKX2.1 HCR probe set. Finally, we would like to thank both Claudio Stern and Marysia Placzek for valuable comments on earlier versions of the work.

## Author Contributions

Conceptualisation: B.S., A.N..; Experimentation: A.N.; Supervision: B.S., Writing: A.N.; Reviewing & editing: A.N., B.S.

