## Supplementary figures for "Dissecting planar and vertical organiser signals in early chick neural development"

Supplementary Information

***Supplementary table 1. Quantification of frequency of neural protrusion formation in anterior segments isolated from the posterior body at different stages of development****.* *The frequency of neural protrusion formation in anterior segments was quantified after isolation at HH2-HH6. Isolation at HH2 led to protrusion formation more frequently (44%) than isolation at HH3 (23.91%). In most cases, protrusions from cultures isolated at HH2 were associated with CHORDIN expression, suggesting that regenerated node tissue contributed to the morphogenesis of the protrusion. Anterior segments isolated from later stage embryos (HH3$^{+}$-HH6) were less frequently associated with CHORDIN expression, meaning that morphogenesis of the neural tissue often occurred in the absence of node tissue.*

| Stage of isolation | Total number of anterior segments cultures | Number of anterior segments that formed protrusions | % formed protrusions |
|---|---|---|---|
| **HH2** | 25 | 11 | 44.00 |
| **HH3** | 46 | 11 | 23.91 |
| **HH3$^{+}$** | 23 | 9 | 39.13 |
| **HH4** | 41 | 35 | 85.37 |
| **HH5** | 16 | 16 | 100.00 |
| **HH6** | 5 | 5 | 100.00 |

*Supplementary table 2: RT-qPCR primer sequences.*

| Gene | Forward primer | Reverse primer |
|---|---|---|
| *ACTB* | TGGCAATGAGAGGTTCAGGT | ATGCCAGGGTACATTGTGGT |
| *Chordin* | GTGCTGGTGTGTGTCAGGAG | CTGTGGCTTCGTGCATCTTT |
| *Goosecoid* | AGACGGCACCGGACTATCTT | CCAAACCTCCACTTTCTCCTC |
| *Lhx1* | TGTGGTTCACTGTGCTAGTTTG | CGGAGCACCTGACAGTATTT |
| *Shh* | TTGGTACTCACGGCTCCTCT | TCCCACACTCTGTCTCTGTCC |

*Supplementary table 3: HCR probes obtained from Molecular Instruments.*

| Gene | NCBI Accession Number | Initiator |
|---|---|---|
| *Chordin* | AF031230.1 | B1 |
| *Goosecoid* | NM_205331.2 | B5 |
| *Lhx1* | NM_001397359.1 | B3 |
| *Shh* | NM_204821.1 | B4 |
| *Otx2* | NM_204520.2 | B2 |
| *En2* | NM_001267719.2 | B5 |
| *Krox-20* | XM_025152050.3 | B3 |
| *Sox3* | NM_204195.2 | B2 |
| *Sox2* | NM_205188.3 | B5 |
| *Sox1* | NM_204333.1 | B4 |
| *Six3* | NM_204364.2 | B2 |

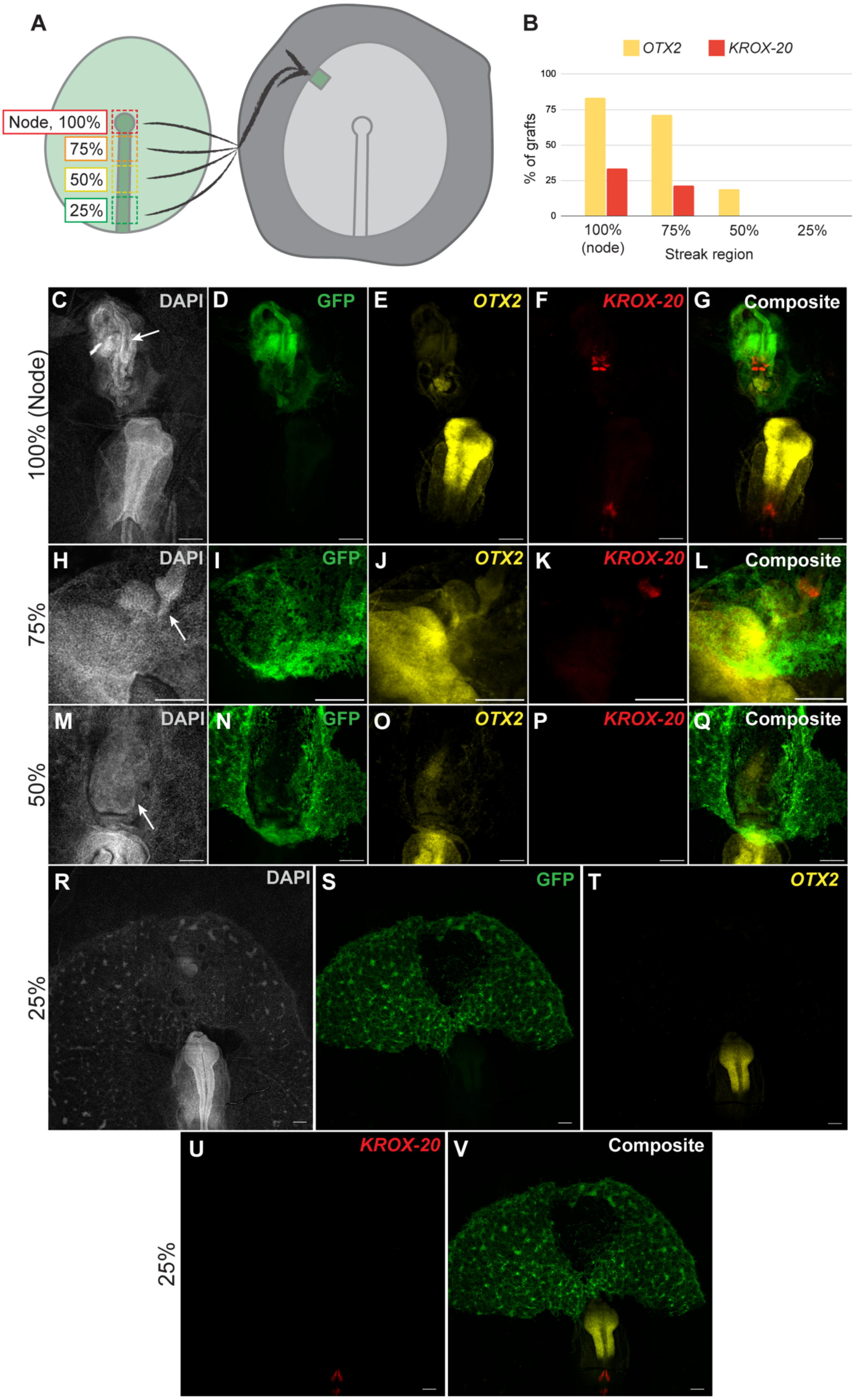
A
Node, 100%
75%
50%
25%
B
OTX2
KROX-20
100
75
50
25
0
% of grafts
100% (node)
75%
50%
25%
Streak region
C
DAPI
D
GFP
E
OTX2
F
KROX-20
G
Composite
100% (Node)
H
DAPI
I
GFP
J
OTX2
K
KROX-20
L
Composite
75%
M
DAPI
N
GFP
O
OTX2
P
KROX-20
Q
Composite
50%
R
DAPI
S
GFP
T
OTX2
25%
U
KROX-20
V
Composite
25%

**Supplementary Figure 1. Neural induction ability is distributed along the primitive streak.** (A) Diagram showing the experimental method; the streak from a $GFP^+$ embryo was divided into 4 sections along the AP axis, and grafted to the outer area pellucida of a wild-type embryo, before culturing for 24h. (B) Quantification of the percentage of grafts from each region inducing *OTX2* (yellow) or *KROX-20* (red). n=24 (node graft), 14 (75% region), 16 (50% region), 3 (25% region). (C-G) After a node graft, the ectopic axis expresses both *OTX2* and *KROX-20*. (H-L) Example of a 75% region graft in which the ectopic axis expresses both *OTX2* and *KROX-20*. (M-Q) Example of a 50% region graft, in which an ectopic axis has not developed, but a patch of ectopic *OTX2* expression is visible. (R-V) shows an example of a 25% region graft, in which there is no ectopic axis or induction of gene expression. All images are 5X stitched maximum intensity projections, scale bars 200µm. Arrows indicate the induced structure.

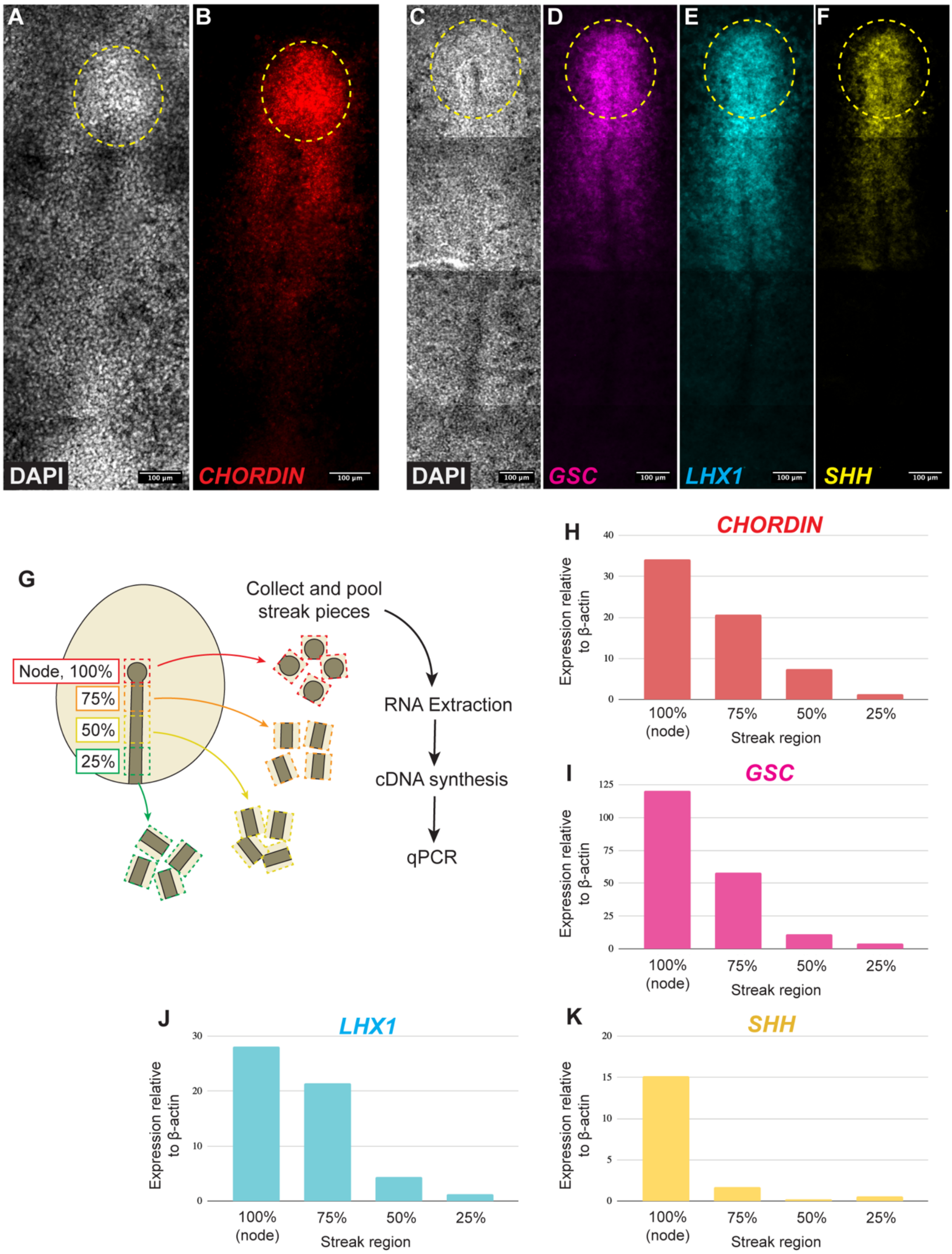


**Supplementary Figure 2. Node marker genes are expressed along the primitive streak.** (A-F) DAPI and HCR FISH staining showing the expression of node marker genes: (B) *CHORDIN*, (D) *GOOSECOID (GSC)*, (E) *LHX1* and (F) *SHH*. The approximate position of the node is denoted by the dotted line. All images are 5X stitched maximum intensity projections, scale bars 100µm. (G) Diagram showing the experimental method for RT-qPCR of sections of the primitive streak from different anteroposterior levels. Primitive streak pieces from 18 $HH3^{+}$ embryos were pooled, before RNA extraction, cDNA synthesis and qPCR. qPCR results are shown for (H), *CHORDIN*, (I) *GOOSECOID (GSC)*, (J) *LHX1* and (K) *SHH*.

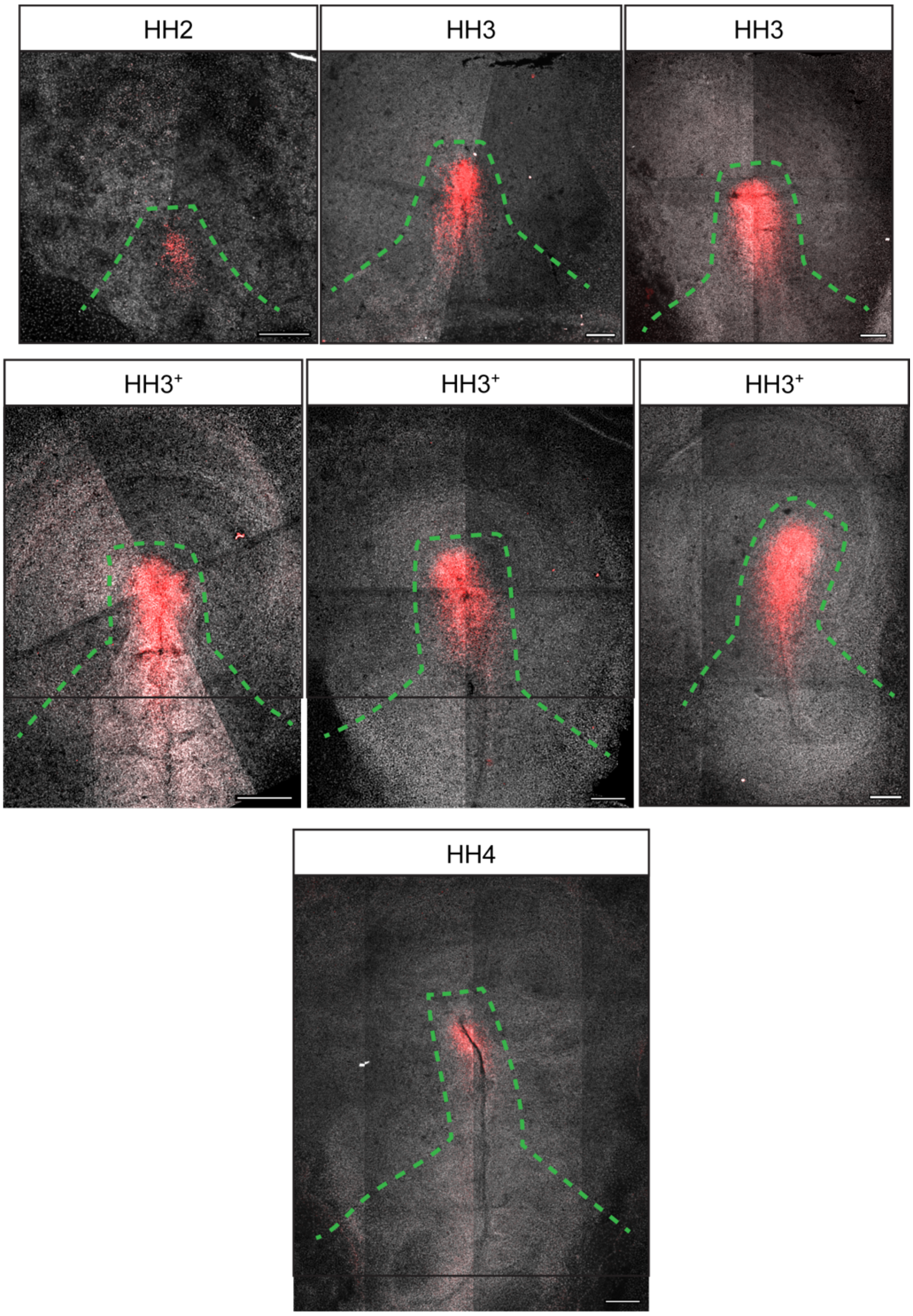


**Supplementary Figure 3. A range of control embryos at different stages stained with DAPI (grey) and HCR-ISH for *CHORDIN* (red).** The dotted lines indicate the approximate locations of cuts performed to isolate the anterior segment. Cutting was carried out with the aim of ensuring that prospective neural tissue from all AP regions of the nervous system would be included, while avoiding the inclusion of *CHORDIN*+ tissue. 5X maximum intensity projections, scale bars 200µm.

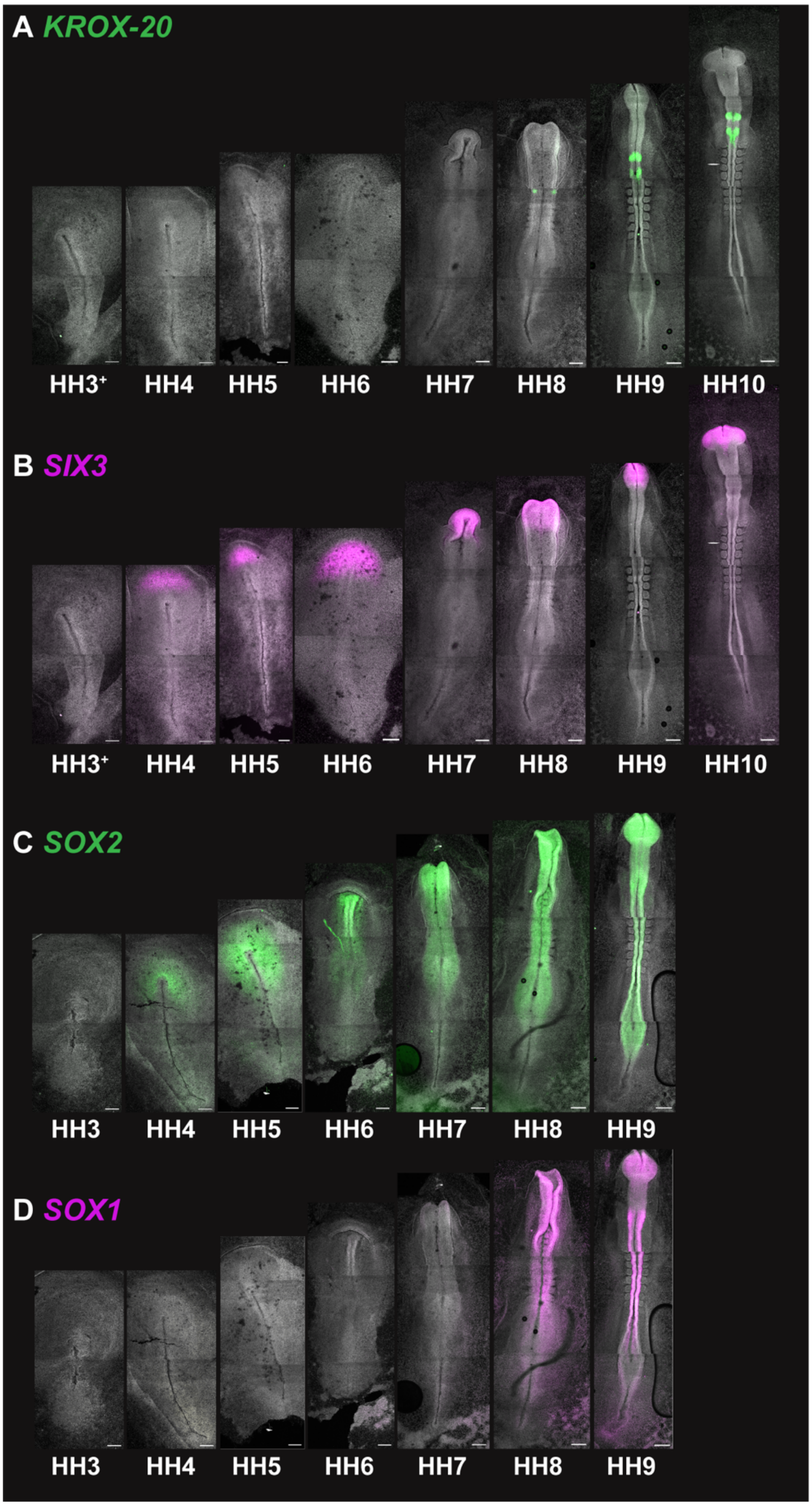


**Supplementary Figure 4. A time course of expression for (A) *KROX-20*, (B) *SIX3*, (D) *SOX2*, and (D) *SOX1*.** All images are 5X maximum intensity projections, scale bars 200µm.
